# 17-Oxime ethers of oxidized ecdysteroid derivatives modulate oxidative stress in human brain endothelial cells and dose-dependently might protect or damage the blood-brain barrier

**DOI:** 10.1101/2023.08.11.552943

**Authors:** Máté Vágvölgyi, Dávid Laczkó, Ana Raquel Santa-Maria, Fruzsina R. Walter, Róbert Berkecz, Mária Deli, Gábor Tóth, Attila Hunyadi

## Abstract

20-Hydroxyecdysone and several of its oxidized derivatives exert cytoprotective effect in mammals including ans. Inspired by this bioactivity of ecdysteroids, in the current study it was our aim to prepare a set of sidechainmodified derivatives and to evaluate their potential to protect the blood-brain barrier (BBB) from oxidative stress. Six novel ecdysteroids, including an oxime and five oxime ethers, were obtained through regioselective synthesis from a sidechain-cleaved calonysterone derivative **2** and fully characterized by comprehensive NMR techniques revealing their complete ^1^H and ^13^C signal assignments. Surprisingly, several compounds sensitized hCMEC/D3 brain microvascular endothelial cells to *tert*-butyl hydroperoxide (tBHP)-induced oxidative damage as recorded by impedance measurements. Compound **8**, containing a benzyloxime ether moiety in its sidechain, was the only one that exerted a protective effect in a higher, 10 μM dose, while at lower (10 μM – 1 μM) doses it promoted tBHP-induced cellular damage. Based on our results, 17-oxime ethers of oxidized ecdysteroids modulate oxidative stress of the BBB in a way that may point towards unexpected toxicity. Further studies are needed to evaluate any possible risk connected to dietary ecdysteroid consumption and CNS pathologies in which BBB damage plays an important role.

## Introduction

Ecdysteroids are insect molting hormone analogs widespread in the Plant Kingdom, and they have attracted a significant interest due to their non-hormonal anabolic, cytoprotective, and vascular protective activity in mammals. Several recent clinical trials have set their focus on these compounds as potential therapeutic agents in the treatment of sarcopenia (NCT03021798, NCT03452488), or the frequently fatal respiratory deterioration in COVID-19 (NCT04472728).

20-Hydroxyecdysone (20E), the most abundant representative of this compound group, was previously found to exert neuroprotective activity in rodent models of cerebral ischemia/reperfusion [1, 2]. Further, we have recently reported a set of three new, highly oxidized ecdysteroids to protect human brain endothelial cells from oxidative injury [3].

The chemical modification of lipids, proteins, and DNA by reactive oxygen species (ROS) can result in cellular and tissue damage, implicating oxidative stress in the pathogenesis of numerous diseases and injuries affecting almost every organ system. Although oxidative stress and its related disorders are more prevalent in older individuals, environmental factors such as air pollution and UV exposure can expedite the development of these conditions in people of all ages [4]. The impact of oxidative stress on neurodegenerative diseases is of significant interest, as it has been linked to the severity of disease pathology. Biomarkers such as peroxiredoxins and ubiquinone/ubiquinol are found to be elevated in individuals with Alzheimer’s disease, Parkinson’s disease, and amyotrophic lateral sclerosis, and are associated with cognitive impairment [4-6]. The blood-brain barrier (BBB) plays a crucial role in ROS-mediated injury and neurodegenerative diseases. This barrier is composed of endothelial cells that have a strong and dynamic interaction with the neighboring cells pericytes and astrocytes. The brain endothelial cells have a unique protection system and it controls the transport of substances in and out of the brain via tight junctions, transport pathways, and efflux proteins [7]. Given that ROS can affect brain endothelial cells and cause BBB disruption, it is crucial to explore whether compounds such as, e.g., ecdysteroids, can provide protection to these cells, promoting BBB protection in the early stages of neurological diseases.

The cytoprotective effect of 20E is at least partly due to its ability to activate protein kinase B (Akt) [8]. In our previous studies on various ecdysteroids as activators of this kinase, we found that calonysterone (**1**), and particularly its side-chain cleaved derivative **2**, are more potent in this regard than 20E [9, 10].

Our previous studies on ecdysteroid oximes and oxime ethers revealed that poststerone, the side-chain cleaved derivative of 20E, can be transformed into 20-oximes and oxime ethers in a regioselective manner [11]. This opened way to a synthetic strategy to prepare ecdysteroid derivatives with a modified, nitrogen-containing side-chain.

Inspired by the neuro- and cerebrovascular protective activity of natural ecdysteroids against ROS, in this work it was our aim to prepare a set of sidechain-cleaved and oxime ether-containing sidechain-modified derivatives of calonysterone (**1**), and to evaluate the compounds’ bioactivity as potential BBB protecting candidates.

## Results and discussion

### Chemistry

#### Oxidative sidechain cleavage

The regioselective oxidative cleavage between the 20,22-diol to eliminate the sterol side chain at the C-17 position of calonysterone **1** was carried out with the hypervalent iodine reagent (diacetoxyiodo)benzene (PIDA), which had been successfully used for the similar purpose in the case of 20-hydroxyecdysone [12]. According to our previous results, using this reagent leads to a significantly better yield than PIFA, a more aggressive oxidant [12]. Full conversion was achieved within an hour. After neutralization and evaporation of the solvent, normal-phase chromatography was used for purification; this was a more practical choice than reverse-phase separation due to its higher loading capacity and milder solvent evaporation conditions. Outline of the reaction is shown in (Fig. 1).

**Fig 1.**
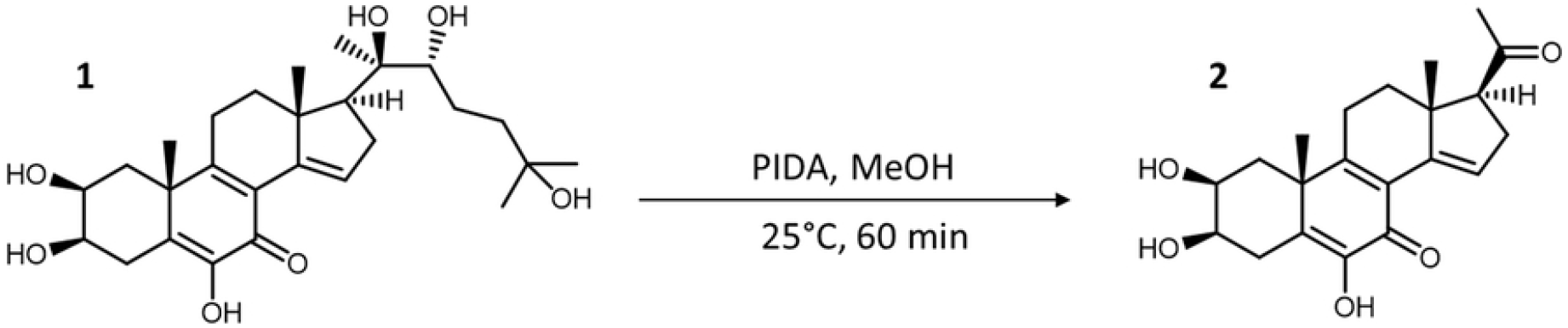
Oxidative cleavage of the sterol sidechain of calonysterone (**1**).

Regioselective formation of 20-oxime or -oxime ether function. Previously, we reported that the 6-enone function of poststerone is relatively less reactive for oxime formation than its 17-oxo group [13], therefore it was postulated that a similar regioselective oximation should be straightforward also for compound **2**.

At first, we performed small-scale (with approx. 10 mg of substrate) test reactions monitored by TLC in every 10 minutes, and we found that all reactions reached full conversion within 40 minutes. Our experiments included the use of either pyridine or ethanol as solvent, and our experience showed that the reactions proceeded in both solvents with nearly identical results. Therefore, we chose ethanol considering its lower boiling point that makes it easier to evaporate during the work-up. After the transformations, the solvent was evaporated on a rotary vacuum evaporator and liquid-liquid extraction was performed with water and ethyl acetate. Outline of the synthesis and structure of the products is shown in (Fig 2).

**Fig 2.**
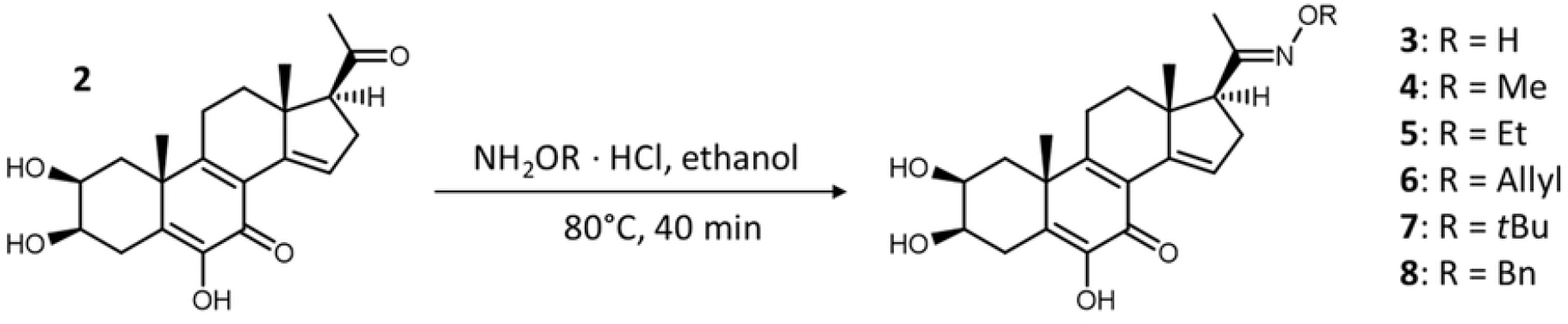
Synthesis of oxime (**3**) and oxime ether (**4–8**) derivatives of compound **2**.

Following this strategy, a total of six new ecdysteroid C-20 oxime and oxime ether derivatives were synthesized from larger-scale aliquots of compound **2**. After pre-purification of the synthesized materials, their HPLC chromatograms were recorded, which was accompanied by the mapping of the eluent systems for their preparative RP-HPLC purification. To improve sample solubility for preparative RP-HPLC, a 3:7 (v/v) ratio solvent mixture of dimethyl sulfoxide (DMSO) and acetonitrile was used.

#### Structure elucidation

We have recently reported the structure elucidation and complete ^1^H and ^13^C signal assignment of compound **1** [9] and the sidechain-cleaved calonysterone derivative **2** [12]. Structure elucidation of the new compounds **3–8** (Fig 2) was performed based on the molecular formulas obtained by HRMS and on detailed NMR studies. HRMS data obtained verified that our synthetic oximation procedure was regioselective in each case, and the reaction took place at either the 6- or 20-carbonyl groups of the substrate. The location and identity of the newly formed functions was determined by means of comprehensive one- and two-dimensional NMR methods using widely accepted strategies [14, 15].

^1^H NMR, ^13^C DeptQ, edHSQC, HMBC, one-dimensional selective ROESY (Rotating frame Overhauser Enchancement Spectroscopy) spectra (τ_mix_: 300ms) were utilized to achieve complete ^1^H and ^13^C signal assignment. It is worth mentioning that due to the molecular mass of compounds **3–8** (374–464 Da) the signal/noise value of the selective ROE experiments strongly exceeds that of the selective NOEs. ^1^H assignments were accomplished using general knowledge of chemical shift dispersion with the aid of the ^1^H-^1^H coupling pattern. ^1^H and ^13^C chemical shifts (600 and 150 MHz, respectively), multiplicities and coupling constants of compounds **3–8** are compiled in (Table 1). Since the stereostructure of the steroid frame is identical within these compounds, we described the multiplicity and *J* coupling constants only for 3. The characteristic NMR (Fig. S1–S20) and HRMS (Fig. S21–S20) spectra of compounds **3–8** are presented as Supporting Information. To facilitate the understanding of the ^1^H and ^13^C signal assignments, the compounds’ structures are also depicted on the spectra.

**Table 1.**
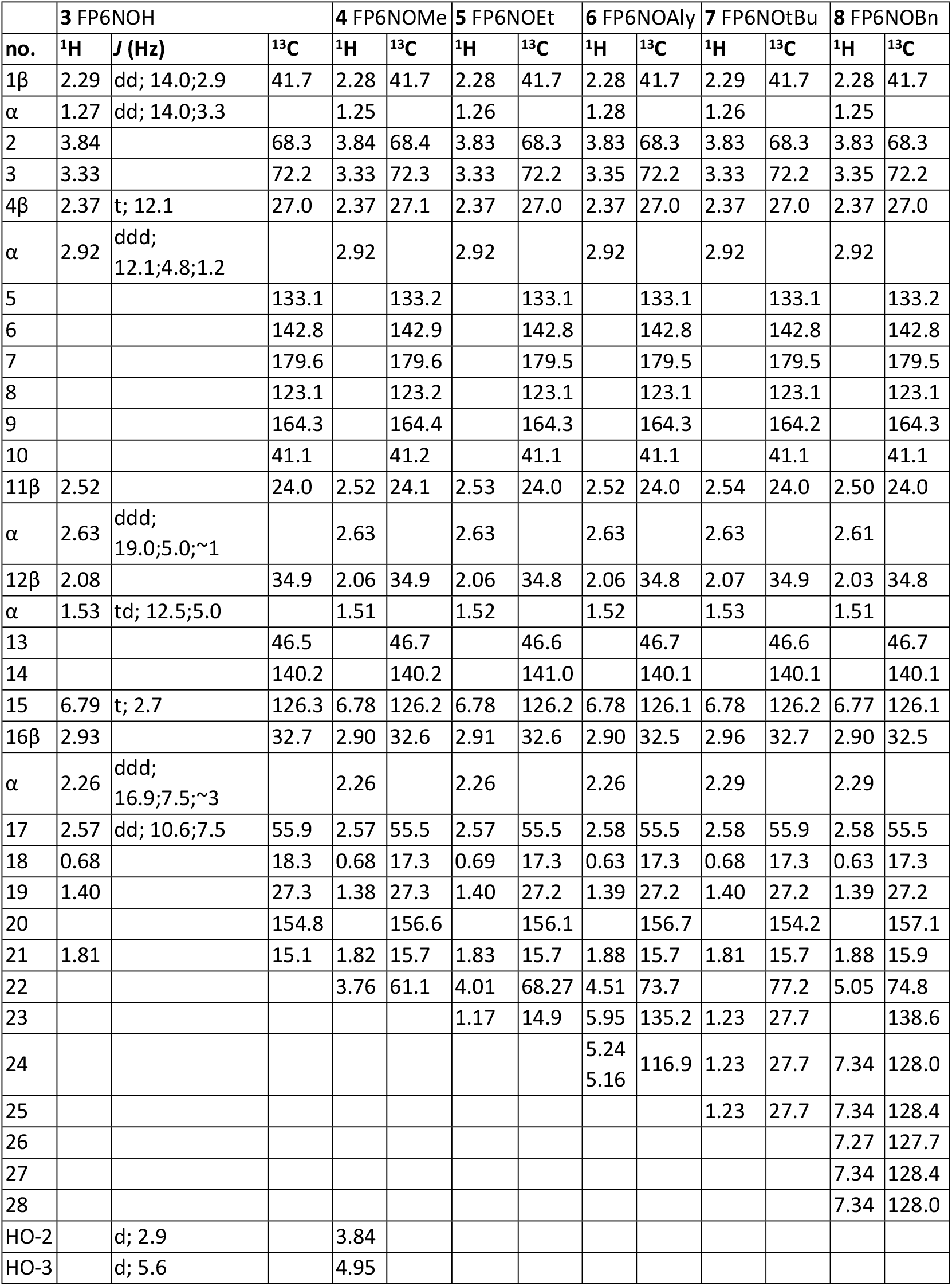
^1^H and ^13^C chemical shifts, multiplicities and coupling constants of compounds **3–8** in dmso-*d*_6._

Only one set of signals appeared in the ^1^H and ^13^C NMR spectra of each compound, indicating that the regioselective oximation led to the isolation of one stereoisomer for each. The measured Δδ 55 ppm diamagnetic change of δC-20 (211 → 156 ppm) supported the C═O → C═NOR conversion [11]. During the NMR study of isomeric Z/E 6-oxime derivatives of 20-hydroxyecdysone 2,3;20,22-diacetonide the chemical shift of α carbon atoms (δC-5 and δC-7) in the syn position with respect to the oxime hydroxyl group exhibits a significant (Δδ syn-anti ∼ 5 ppm) diamagnetic shift, which was successfully utilized for differentiation of (Z/E) isomers [16]. In the present case for compounds **3**–**8**, due to the absence of exact data of Δδ syn-anti parameters for the C-21 and C-17 signals, the unambiguous identification of the E/Z isomerism in this way was not possible. To overcome this problem, we utilized a series of selective ROESY experiments on the CH_3_-21 signals (Fig. S1, S8 and S13), and the detected steric responses unequivocally proved the *E* configuration of the oxime moiety. By introducing the 1D selROE spectrum on H_3_-21 into the edited HSQC experiment (Fig. S3, S8 and S13), the ROE signals allowed identifying the corresponding C−H cross-peaks. The quaternary carbon signals were identified from the HMBC spectra, for which the HMBC responses over two and three bonds of H_3_-19, H_3_-18, and H_3_-21 were very effective (Fig S4, S9 and S14).

#### Biology

We evaluated the effect of the compounds on the viability and barrier integrity of human brain microvascular endothelial cells (hCMEC/D3) using impedance measurements. Initially, we tested concentration ranges of 0.01–10 µM for all compounds, and no notable changes in cell viability were observed, except for compounds **3, 4** and **8** (Supporting Information, Fig S27). Although we monitored all concentrations for 24 hours, we observed that the effect of all compounds starts occurring at the 4-hour time point. Therefore, we have focused our results on the 4h timepoint. For compound **3** we could observe a significant cell index decrease for 10 µM concentration, however, for compound **4** a significant increase for 1 µM concentration was observed (Supporting Information, Fig S27). Notably, compound **8** exhibited the highest and most significant activity. At concentrations of 0.01, 0.1, 1, and 10 µM, it demonstrated a positive effect on barrier integrity. As compound **8** demonstrated the highest activity, we decided to investigate whether it also promotes a protective effect against oxidative stress. Excessive ROS resulting from oxidative stress can cause disruption of the BBB by compromising the antioxidant defense system. The damaging effects of ROS on cellular components such as proteins, lipids, and DNA can lead to the modulation of tight junctions, activation of matrix metalloproteinases, and upregulation of inflammatory molecules, all of which can contribute to BBB damage [17]. Tert-butyl hydroperoxide is known to induce cellular damage by generating high levels of ROS [18]. Therefore, to assess the protective effects of the compound against ROS-induced damage, we treated cells with tBHP (350 µM) alone or in combination with 0.01, 0.1, 1 and 10 µM of compound **8**. We identified the optimal concentration of tBHP by testing different concentrations and selecting 350 µM, which did not decrease the cell index below ∼50% in our previous work. This concentration was then used for the cell viability assay.

We can observe a significant decrease in cell viability by a total of ca. 60% in the presence of tBHP compared to the control group (Fig 3B–C), indicating tBHP-induced oxidative damage on the cells. Treatment of 10 µM of compound **8** resulted in a significant and steady increase of cell impedance, i.e., it was able to protect the cells efficiently from the harmful effects of tBHP. These findings suggest that the compound might have a protective effect against cellular damage induced by ROS (Fig 3B and 3C). However, at smaller, 10 nM, 100 nM, and 1 μM concentrations a surprising opposite effect was observed. In the lower concentrations and with at least 6h incubation, compound **8** dramatically increased tBHP-induced toxicity, leading to a nearly complete disruption of the cellular layer (Fig 3C). We also tested compounds **4** and **6** (3 and 10 µM) in combination with 350 µM of tBHP, and both significantly increased oxidative damage at these concentrations (Supporting Information, Fig S28).

**Fig 3.**
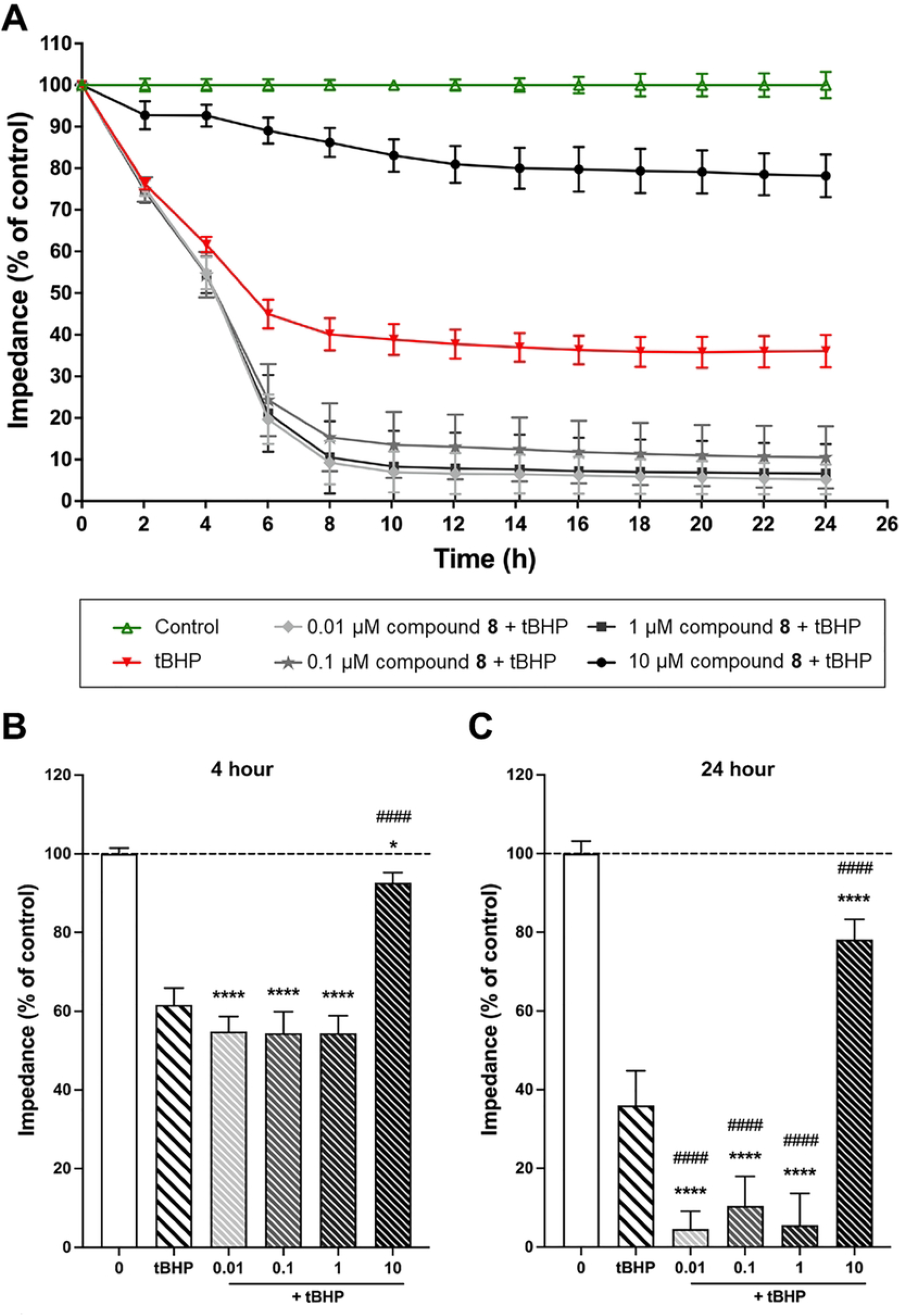
The effects of compound **8** at concentrations of 0.01, 0.1, 1, and 10 μM treatment on human brain microvascular endothelial cells (hCMEC/D3) were evaluated using impedance-based assays to assess cell viability and barrier integrity in the absence and presence of oxidative stress promoted by tert-butyl hydroperoxide (tBHP). **A**: Time-dependent impact of **8** on cell viability following cotreatment with tBHP (350 µM). **B**: Impact of **8** on cell viability at 4 hours co-treatment with tBHP (350 µM). **C**: Impact of **8** on cell viability at 24 hours co-treatment with tBHP (350 µM). The data are presented as the mean ± standard deviation (SD) and were obtained from a minimum of two independent experiments (n = 2−3) with 3−9 technical replicates. Data analysis was performed using one-way analysis of variance (ANOVA) followed by Dunnett’s multiple comparisons test. The results were statistically significant with *p < 0.05, ****p < 0.0001, compared to the control group, and ^####^p < 0.0001, compared to the tBHP group.

The use of impedance-based monitoring to assess brain endothelial cell function is crucial as it not only measures the number of viable cells but also provides valuable information on the integrity of the cell layer and the extent of barrier damage. This method has been shown to be relevant to evaluate barrier integrity and the overall health of brain endothelial cells [3, 19, 20]. There is evidence indicating that oxidative stress plays a crucial role in the induction of BBB damage [7]. The present study provided evidence that treatment with tBHP resulted in brain endothelial damage, which was manifested by a decrease in cell and barrier integrity in certain concentrations of the compounds tested. However, co-treatment with compound **8** significantly altered this effect, leading to the prevention or promotion of oxidative barrier damage. During the 24h-long monitoring of the cell index, a clear concentration-dependent distinction could be made between the protective or damaging effect. To this end, no data are available on the pharmacokinetics of compound **8**, hence it is not possible to evaluate if a 10 µM plasma level at the BBB is achievable or not. On the other hand, the low-concentration effect of compound **8** to sensitize the BBB to oxidative stress clearly raises a warning concerning its value as a lead compound.

In the broader context, it may be worth stressing that the herein reported compounds are semi-synthetic ecdysteroids that contain oxime ether moieties in their sidechain. This functional group is not expectable to occur in natural ecdysteroids or their metabolites, therefore our results do not directly imply any risk connected to phytoecdysteroid consumption. In our previous study on minor phytoecdysteroids [3], only protective effects were observed. Nevertheless, considering that hardly anything is known about ecdysteroids’ bioactivity in relation with the BBB, further studies are needed to evaluate related drug discovery potential and/or risks of this compound family.

### Experimental Materials and methods

#### Chemistry

All solvents and reagents were purchased from Sigma-Aldrich (Merck KGaA, Darmstadt, Germany) and were used without any further purification. The progress of the reactions was monitored by thin layer chromatography (TLC) on Kieselgel 60F254 silica plates purchased from Merck (Merck KGaA, Darmstadt, Germany). The examination of the plates was carried out under UV illumination at 254 and 366 nm.

The purification of calonysterone was performed by centrifugal partition chromatography on a 250 ml Armen Spot instrument (Gilson Inc., Middleton, WI, USA). The flash chromatographic purification of compound **2** was carried out on a Combiflash Rf+ instrument (Teledyne ISCO, Lincoln, NE, USA) equipped with diode array and evaporative light scattering detection (DAD-ELSD), and commercially available prefilled RediSep columns (Teledyne ISCO, Lincoln, NE, USA) were utilized. For the analysis of the compounds, we used a dual-pump Jasco HPLC instrument (Jasco International Co. Ltd., Hachioji, Tokyo, Japan) equipped with an “MD-2010 Plus” PDA detector. The analytical-scale separations were performed on a Phenomenex Kinetex Biphenyl 100A 5µ 250x4.6 mm (Torrence, CA, USA) HPLC column. The separation of compound 4–8 was performed on an Armen “Spot Prep II” preparative chromatographic apparatus (Gilson Inc., Middleton, WI, USA) equipped with a dual-wavelength UV detector and four individual solvent pumps. The RP-HPLC purification of the ecdysteroid products was carried out with adequately chosen isocratic eluent mixtures of acetonitrile and water.

#### Isolation of calonysterone (1)

A commercially available extract prepared from *Cyanotis arachnoidea* roots was purchased from Xi’an Olin Biological Technology Co., Ltd. (Xi’an, China) [21], and subjected to a chromatographic purification to obtain the starting material calonysterone (**1**) as published before [22]. Briefly, 5.46 kg of extract was percolated with 15.5 L of methanol, and after evaporation of the solvent, the dry residue (700 g) was subjected to further separation by a multi-step chromatographic fractionation through silica gel. The final purification of **1** was carried out using centrifugal partition chromatography in ascending mode with a biphasic solvent system of *n*-hexane – ethyl acetate – methanol – water (1:5:1:5, v/v/v/v) [22].

#### Preparation of compound 2 via the oxidative side-chain cleavage of calonysterone (1)

An aliquot of 2 g of calonysterone (**1**) was dissolved in 500 ml of methanol. One equivalent (1.34 g) of PIDA was added, and the reaction mixture was stirred for 60 minutes at room temperature. The solution was then neutralized with 10% aq. NaHCO_3_, and the solvent was evaporated under reduced pressure on a rotary evaporator. Subsequently, the residue was re-dissolved in acetone and adsorbed on 10 g of silica gel for dry loading. The product was purified by flash chromatography on a 24 g silica column (flow rate 35 ml/min, run time: 60 min) with a gradient of dichloromethane (A) and methanol (B), from 0% to 15% of solvent B in A. The separation afforded compound **2** in a yield of 45.5%.

#### General Procedure for the synthesis of sidechain cleaved calonysterone 20-oxime and oxime ether derivatives 3–8

A 120 mg aliquot of compound **2** (0,34 mmol) was dissolved in ethanol (20 ml) and depending on the functional group to be coupled, 120 mg of hydroxylamine hydrochloride (compound **3**) or alkoxyamine hydrochloride (compounds **4**–**8**) was added to the solution under stirring. After 40 minutes of stirring at 80°C the solution was evaporated to dryness under reduced pressure. After water addition to the dry residue (100 ml), the aqueous solution was extracted three times with ethyl acetate (3x100 ml) and the combined organic phase was dried over anhydrous Na_2_SO_4_. Subsequently, the solution was filtered, and the solvent was evaporated under reduced pressure. The purification of the mixture was implemented by preparative RP-HPLC to afford the corresponding ecdysteroid product.

#### Procedures for structure elucidation of the obtained products

HR-MS analysis of the compounds was carried out on an Agilent 1100 LC-MS instrument (Agilent Technologies, Santa Clara, CA, USA) coupled with Thermo Q-Exactive Plus orbitrap spectrometer (Thermo Fisher Scientific, Waltham, MA, USA) used in positive ionization mode. Regarding the samples, 100 µg/ml solutions were prepared with acetonitrile solvent containing 0.1% formic acid.

^1^H NMR, ^13^C DeptQ, edHSQC, HMBC, and one-dimensional selective ROESY spectra (τ_mix_: 300ms) were recorded at 295 K on a Bruker Avance III HD 600 (Billerica, MA, USA; 600 and 150 MHz for ^1^H and ^13^C NMR spectra, respectively) spectrometer equipped with a Prodigy cryo-probehead. The pulse programs were taken from the Bruker software library (TopSpin 3.5). ^1^H assignments were accomplished using general knowledge of chemical shift dispersion with the aid of the ^1^H-^1^H coupling pattern (^1^H NMR spectra). DMSO-*d*_6_ were used as the solvent and tetramethylsilane (TMS) as the internal standard and amounts of approximately 1–5 mg of compound was dissolved in 0.1 ml of solvent and transferred to 2.5 mm Bruker MATCH NMR sample tube (Bruker). Chemical shifts (δ) and coupling constants (*J*) are given in ppm and in Hz, respectively. To facilitate the understanding of the ^1^H and ^13^C signal assignments, the structures are also depicted on the spectra (Supporting Information, Fig S1–S20).

**Compound 3**: off-white, solid; isolated yield: 49.4 mg (39.5%); RP-HPLC purity: 98.1%; for ^1^H and ^13^C NMR data, see Table 1 and Supplementary Fig. S1–S4; HR-MS: C_21_H_27_NO_5_, [M+H]^+^ Calcd.: 374.19730, found: 374.19696 (Fig S21).

**Compound 4**: off-white, solid; isolated yield: 33.1 mg (25.5%); RP-HPLC purity: 99.1%; for ^1^H and ^13^C NMR data, see Table 1 and Supplementary Fig. S5–S9; HR-MS: C_22_H_29_NO_5_, [M+H]^+^ Calcd.: 388.21185, found: 388.21208 (Fig S22).

**Compound 5**: off-white, solid; isolated yield: 50.6 mg (37.7%); RP-HPLC purity: 98.0%; for ^1^H and ^13^C NMR data, see Table 1, and Supplementary Fig. S10–S14; HR-MS: C_23_H_31_NO_5_, [M+H]^+^ Calcd.: 402.22750, found: 402.22795 (Fig S23).

**Compound 6**: off-white, solid; isolated yield: 52.1 mg (37.7%); RP-HPLC purity: 98.9%; for ^1^H and ^13^C NMR data, see Table 1, and Supplementary Fig. S15–S16; HR-MS: C_24_H_31_NO_5_, [M+H]+ Calcd.: 414.22750, found: 414.22808 (Fig S24).

**Compound 7**: off-white, solid; isolated yield: 27.8 mg (19.4%); RP-HPLC purity: 97.4%; for ^1^H and ^13^C NMR data, see Table 1, and Supplementary Fig. S17–S18; HR-MS: C_25_H_35_NO_5_, [M+H]^+^ Calcd.: 430.25880, found: 430.25890 (Fig S25).

**Compound 8**: off-white, solid; isolated yield: 54.9 mg (35.3%); RP-HPLC purity: 97.1%; for ^1^H and ^13^C NMR data, see Table 1, and Supplementary Fig. S19–S20; HR-MS: C_28_H_33_NO_5_, [M+H]^+^ Calcd.: 464.24315, found: 464.24351 (Fig S26).

### Biology

#### Human brain microvascular endothelial cell line (hCMEC/D3) as a blood-brain barrier cell culture model

The hCMEC/D3 human brain microvascular endothelial cell line was obtained from Merck Millipore (Germany). To maintain the cells’ brain endothelial-like features, we used cells under passage number 35 [23]. Cells were grown in dishes coated with rat tail collagen and maintained in an incubator at 37°C with 5% CO_2_. The basal medium used was MCDB 131 (Pan Biotech, Germany) supplemented with 5% fetal bovine serum, GlutaMAX (100 ×, Life Technologies, USA), lipid supplement (100 ×, Life Technologies, USA), 10 µg/ml ascorbic acid, 550 nM hydrocortisone, 37.5 µg/ml heparin, 1 ng/ml basic fibroblast growth factor (Roche, USA), 5 µg/ml insulin, 5 µg/ml transferrin, 5 ng/ml selenium supplement (100x, PanBiotech), 10 mM HEPES, and gentamycin (50 µg/ml). We changed the medium every two or three days. When the cultures reached almost 90% confluence, we passaged them to rat tail collagen-coated 96-well plates (E-plate, Agilent, USA) for viability assays. Before each experiment, the medium was supplemented with 10 mM LiCl for 24 hours to improve BBB properties [24].

#### Impedance measurements for cell viability assays

The viability of brain endothelial cells was assessed using real-time impedance measurement, which has been shown to correlate with cell number, adherence, growth, and viability [25]. The hCMEC/D3 cells were seeded in 96-well E-plates with golden electrodes at a density of 5×10^3^ cells per well and incubated in a CO_2_ incubator at 37°C for 5-6 days. The medium was changed every two days. Once the cells reached a stable growth plateau, they were treated with compounds **2**–**8** at concentrations ranging from 0.01 to 10µM, and their viability was monitored for 24 hours using RTCA-SP (Agilent). Triton X-100 was used to determine 100% toxicity. After 24-hours monitoring, we found that the compounds exhibited the highest level of activity after 4-hours of treatment. As a result, we decided to focus our treatments at this time point.

#### Preparation of stock and working solutions for the cellular assays

The compounds were obtained as dry powder and stored at -20 °C until use. Stock solutions were prepared by diluting the compounds in DMSO to a final concentration of 10 mM and stored at -20 °C. Working solutions were freshly prepared by diluting the stock solutions in cell culture medium to obtain a concentration range of 0.01– 10 µM.

#### Induction of oxidative stress by *tert*-butyl hydroperoxide

The oxidative compound tert-butyl hydroperoxide (tBHP) can cause cell death through apoptosis or necrosis by generating tert-butoxyl radicals via iron-dependent reactions. This results in lipid peroxidation, depletion of intracellular glutathione, and modification of protein thiols, leading to loss of cell viability [18, 26, 27]. To determine a concentration that would result in approximately 50% cell viability loss, various concentrations of tBHP were tested ranging from 1–1000 µM in preliminary experiments [3]. Based on these results, 350 µM tBHP was found to be effective and was used in combination with the selected concentrations of the compounds to test for potential protective effects.

### Statistics

The mean ± SD values were used to present the data. The statistical significance between different treatment groups was assessed using one-way ANOVA, followed by Dunnett’s multiple comparison post-tests (GraphPad Prism 9.0; GraphPad Software, USA). At least four parallel samples were used, and changes were considered statistically significant when p < 0.05.

## Conclusions

In this study, we have prepared a sidechain cleaved, oxidized ecdysteroid and six of its oxime or oxime ether derivatives. Using a relevant *in vitro* cellular model for blood-brain barrier integrity, we demonstrated that the compounds have a significant impact on the oxidative stress-resistance of the BBB. At low doses, compound **8** increased t-BHP-induced cellular damage while at a higher concentration it acted as a protective agent. Our results raise a warning that semi-synthetic modifications of cytoprotective ecdysteroids may unexpectedly alter their bioactivity profile towards harmful effects on cerebrovascular endothelial cells, which may confer them a central nervous system toxicity. The significance of these findings concerning phytoecdysteroid consumption is yet unclear and requires further studies.

## Author Contributions

Conceptualization: A.H., Data curation: A.R.S.M., F.R.W., G.T., Funding acquisition: A.H., F.R.W., M.D., Investigation: M.V., D.L., A.R.S.M., F.R.W., R.B., G.T., Resources: M.D., G.T., A.H., Supervision: M.D., A.H., Writing – original draft: M.V., D.L., G.T., A.H., writing – review & editing: D.L., A.R.S.M., F.R.W., M.D., G.T., A.H.

## References

1. Hu J, Luo CX, Chu WH, Shan YA, Qian Z-M, Zhu G, et al. 20-Hydroxyecdysone Protects against Oxidative Stress-Induced Neuronal Injury by Scavenging Free Radicals and Modulating NF-κB and JNK Pathways. PLOS ONE. 2012;7(12):e50764. doi: 10.1371/journal.pone.0050764.

2. Wang W, Wang T, Feng WY, Wang ZY, Cheng MS, Wang YJ. Ecdysterone protects gerbil brain from temporal global cerebral ischemia/reperfusion injury via preventing neuron apoptosis and deactivating astrocytes and microglia cells. Neurosci Res. 2014;81-82:21–9. Epub 20140127. doi: 10.1016/j.neures.2014.01.005. PubMed PMID: 24480536.

3. Tóth G, Santa-Maria AR, Herke I, Gáti T, Galvis-Montes D, Walter FR, et al. Highly Oxidized Ecdysteroids from a Commercial Cyanotis arachnoidea Root Extract as Potent Blood–Brain Barrier Protective Agents. Journal of Natural Products. 2023;86(4):1074–80. doi: 10.1021/acs.jnatprod.2c00948.

4. Chung TD, Linville RM, Guo Z, Ye R, Jha R, Grifno GN, et al. Effects of acute and chronic oxidative stress on the blood-brain barrier in 2D and 3D in vitro models. Fluids Barriers CNS. 2022;19(1):33. Epub 20220512. doi: 10.1186/s12987-022-00327-x. PubMed PMID: 35551622; PubMed Central PMCID: PMCPMC9097350.

5. Luceri C, Bigagli E, Femia AP, Caderni G, Giovannelli L, Lodovici M. Aging related changes in circulating reactive oxygen species (ROS) and protein carbonyls are indicative of liver oxidative injury. Toxicol Rep. 2018;5:141–5. Epub 20171221. doi: 10.1016/j.toxrep.2017.12.017. PubMed PMID: 29854585; PubMed Central PMCID: PMCPMC5977162.

6. Liguori I, Russo G, Curcio F, Bulli G, Aran L, Della-Morte D, et al. Oxidative stress, aging, and diseases. Clin Interv Aging. 2018;13:757–72. Epub 20180426. doi: 10.2147/CIA.S158513. PubMed PMID: 29731617; PubMed Central PMCID: PMCPMC5927356.

7. Song K, Li Y, Zhang H, An N, Wei Y, Wang L, et al. Oxidative Stress-Mediated Blood-Brain Barrier (BBB) Disruption in Neurological Diseases. Oxidative Medicine and Cellular Longevity. 2020;2020:4356386. doi: 10.1155/2020/4356386.

8. Gorelick-Feldman J, Cohick W, Raskin I. Ecdysteroids elicit a rapid Ca2+ flux leading to Akt activation and increased protein synthesis in skeletal muscle cells. Steroids. 2010;75(10):632–7. Epub 20100402. doi: 10.1016/j.steroids.2010.03.008. PubMed PMID: 20363237; PubMed Central PMCID: PMCPMC3815456.

9. Csábi J, Hsieh TJ, Hasanpour F, Martins A, Kele Z, Gáti T, et al. Oxidized Metabolites of 20-Hydroxyecdysone and Their Activity on Skeletal Muscle Cells: Preparation of a Pair of Desmotropes with Opposite Bioactivities. J Nat Prod. 2015;78(10):2339–45. Epub 20151014. doi: 10.1021/acs.jnatprod.5b00249. PubMed PMID: 26465254.

10. Issaadi HM, Csábi J, Hsieh TJ, Gáti T, Tóth G, Hunyadi A. Side-chain cleaved phytoecdysteroid metabolites as activators of protein kinase B. Bioorg Chem. 2019;82:405–13. Epub 20181031. doi: 10.1016/j.bioorg.2018.10.049. PubMed PMID: 30428419.

11. Bogdán D, Haessner R, Vágvölgyi M, Passarella D, Hunyadi A, Gáti T, et al. Stereochemistry and complete 1H and 13C NMR signal assignment of C-20-oxime derivatives of posterone 2,3-acetonide in solution state. Magnetic Resonance in Chemistry. 2018;56(9):859–66. doi: https://doi.org/10.1002/mrc.4750.

12. Issaadi HM, Csabi J, Hsieh TJ, Gati T, Toth G, Hunyadi A. Side-chain cleaved phytoecdysteroid metabolites as activators of protein kinase B. Bioorg Chem. 2019;82:405–13. Epub 2018/11/15. doi: 10.1016/j.bioorg.2018.10.049. PubMed PMID: 30428419.

13. Vagvolgyi M, Martins A, Kulmany A, Zupko I, Gati T, Simon A, et al. Nitrogen-containing ecdysteroid derivatives vs. multi-drug resistance in cancer: Preparation and antitumor activity of oximes, oxime ethers and a lactam. Eur J Med Chem. 2018;144:730–9. Epub 2018/01/02. doi: 10.1016/j.ejmech.2017.12.032. PubMed PMID: 29291440.

14. Duddeck H, Dietrich W, Tóth G. Structure Elucidation by Modern NMR1998.

15. Pretsch E, Tóth G, Munk EM, Badertscher M. Computer-Aided Structure Elucidation: Wiley-VCH; 2002.

16. Vágvölgyi M, Martins A, Kulmány Á, Zupkó I, Gáti T, Simon A, et al. Nitrogen-containing ecdysteroid derivatives vs. multi-drug resistance in cancer: Preparation and antitumor activity of oximes, oxime ethers and a lactam. Eur J Med Chem. 2018;144:730–9. Epub 20171212. doi: 10.1016/j.ejmech.2017.12.032. PubMed PMID: 29291440.

17. Obermeier B, Daneman R, Ransohoff RM. Development, maintenance and disruption of the blood-brain barrier. Nat Med. 2013;19(12):1584–96. Epub 20131205. doi: 10.1038/nm.3407. PubMed PMID: 24309662; PubMed Central PMCID: PMCPMC4080800.

18. Kucera O, Endlicher R, Rousar T, Lotkova H, Garnol T, Drahota Z, et al. The effect of tert-butyl hydroperoxide-induced oxidative stress on lean and steatotic rat hepatocytes in vitro. Oxid Med Cell Longev. 2014;2014:752506. Epub 20140331. doi: 10.1155/2014/752506. PubMed PMID: 24847414; PubMed Central PMCID: PMCPMC4009166.

19. Harazin A, Bocsik A, Barna L, Kincses A, Varadi J, Fenyvesi F, et al. Protection of cultured brain endothelial cells from cytokine-induced damage by alpha-melanocyte stimulating hormone. PeerJ. 2018;6:e4774. Epub 20180515. doi: 10.7717/peerj.4774. PubMed PMID: 29780671; PubMed Central PMCID: PMCPMC5958884.

20. Santa-Maria AR, Walter FR, Valkai S, Bras AR, Meszaros M, Kincses A, et al. Lidocaine turns the surface charge of biological membranes more positive and changes the permeability of blood-brain barrier culture models. Biochim Biophys Acta Biomembr. 2019;1861(9):1579–91. Epub 20190710. doi: 10.1016/j.bbamem.2019.07.008. PubMed PMID: 31301276.

21. Hunyadi A, Herke I, Lengyel K, Bathori M, Kele Z, Simon A, et al. Ecdysteroid-containing food supplements from Cyanotis arachnoidea on the European market: evidence for spinach product counterfeiting. Sci Rep. 2016;6:37322. Epub 2016/12/09. doi: 10.1038/srep37322. PubMed PMID: 27929032; PubMed Central PMCID: PMCPMC5144001.

22. Issaadi HM, Tsai Y-C, Chang F-R, Hunyadi A. Centrifugal partition chromatography in the isolation of minor ecdysteroids from Cyanotis arachnoidea. Journal of Chromatography B. 2017;1054:44–9. doi: https://doi.org/10.1016/j.jchromb.2017.03.043.

23. Weksler BB, Subileau EA, Perriere N, Charneau P, Holloway K, Leveque M, et al. Blood-brain barrier-specific properties of a human adult brain endothelial cell line. FASEB J. 2005;19(13):1872–4. Epub 20050901. doi: 10.1096/fj.04-3458fje. PubMed PMID: 16141364.

24. Veszelka S, Toth A, Walter FR, Toth AE, Grof I, Meszaros M, et al. Comparison of a Rat Primary Cell-Based Blood-Brain Barrier Model With Epithelial and Brain Endothelial Cell Lines: Gene Expression and Drug Transport. Front Mol Neurosci. 2018;11:166. Epub 20180522. doi: 10.3389/fnmol.2018.00166. PubMed PMID: 29872378; PubMed Central PMCID: PMCPMC5972182.

25. Walter FR, Veszelka S, Pasztoi M, Peterfi ZA, Toth A, Rakhely G, et al. Tesmilifene modifies brain endothelial functions and opens the blood-brain/blood-glioma barrier. J Neurochem. 2015;134(6):1040–54. Epub 20150723. doi: 10.1111/jnc.13207. PubMed PMID: 26112237.

26. Martin C, Martinez R, Navarro R, Ruiz-Sanz JI, Lacort M, Ruiz-Larrea MB. tert-Butyl hydroperoxide-induced lipid signaling in hepatocytes: involvement of glutathione and free radicals. Biochem Pharmacol. 2001;62(6):705–12. doi: 10.1016/s0006-2952(01)00704-3. PubMed PMID: 11551515.

27. Zhao W, Feng H, Sun W, Liu K, Lu JJ, Chen X. Tert-butyl hydroperoxide (t-BHP) induced apoptosis and necroptosis in endothelial cells: Roles of NOX4 and mitochondrion. Redox Biol. 2017;11:524–34. Epub 20170105. doi: 10.1016/j.redox.2016.12.036. PubMed PMID: 28088644; PubMed Central PMCID: PMCPMC5237803.

